# Phosphovariants of the canonical heterotrimeric Gα protein, GPA1, differentially affect G protein activity and Arabidopsis development

**DOI:** 10.64898/2026.01.10.698825

**Authors:** David Chakravorty, Sarah M. Assmann

**Affiliations:** Biology Department, Pennsylvania State University, University Park, Pennsylvania 16802

## Abstract

Heterotrimeric G protein signaling downstream of receptor-like kinases (RLKs) is an emerging and important signal transduction mechanism in plants. However, little is known about the effects of phosphorylation events on the function of the canonical Gα subunit, GPA1. That several known phosphosites reside within important nucleotide co-factor binding sites suggests a role of phosphorylation in modulating the GTP/GDP binding and hydrolysis cycle of GPA1. To mimic and assess the effects of GPA1 phosphorylation, we created ten different phosphovariants of GPA1 and then comprehensively measured *in vitro* biochemical activity, *in vivo* protein-protein interactions, and developmental phenotypes of phosphomutant-complemented Arabidopsis *gpa1* null mutants. Our assays confirmed both that phosphovariants of S49 and S52 in the G1 nucleotide binding motif impair GTP and GDP binding, and that the *gpa1* morphological phenotypes of reduced etiolated hypocotyl elongation, rounded leaves, and short round flowers are not dependent on the nucleotide status of GPA1. Phosphomimetic mutations at S49, S52, T53, and T164 each exhibited a distinct pattern in complementing *gpa1* phenotypes, indicating that GPA1 employs multiple signaling states based on phosphorylation status. The multi-state hypothesis provides a key insight into the mechanism by which a limited repertoire of plant heterotrimeric G protein subunits can transduce a wide variety of signals with exquisite specificity.

## Introduction

Herterotrimeric G proteins represent a critically important class of eukaryotic signal transduction components [1]. The G protein heterotrimer is composed of Gα, Gβ and Gγ subunits. The genome of the model plant Arabidopsis encodes one canonical Gα subunit, GPA1 [2], three extra-large Gα subunits, XLG1, XLG2 and XLG3 [3, 4], one Gβ subunit, AGB1 [5], two canonical Gγ subunits, AGG1 [6] and AGG2 [7], and one extra-large Gγ subunit, AGG3 [8]. In the canonical signaling mechanism, the GDP-bound inactive Gα subunit associates with the Gβγ dimer at the intracellular surface of the plasma membrane. Ligand stimulation of 7-transmembrane (7TM) spanning G protein-coupled receptors (GPCRs) evokes a conformational change in the Gα subunit, resulting in the exchange of GDP for GTP and dissociation of Gα-GTP from the Gβγ dimer. Both Gα-GTP and free Gβγ signal to intracellular effectors until intrinsic GTPase activity of the Gα, which can also be stimulated by regulator of G protein signaling (RGS) proteins, catalyzes the hydrolysis of GTP to GDP, and the return of the complex to the inactive state [9, 10]. However, this simplified paradigm belies substantial layers of complexity identified in G protein signaling. In mammalian systems signaling has been observed from non-dissociated heterotrimers [11] and complexes localized to Golgi and pre-Golgi structures [12, 13]. Furthermore, interdomain motion within the Gα subunit [14], receptor desensitization [15], agonist-antagonist dosage [16] and attenuation of activity by GPCR-mediated GTP release [17, 18] are all proposed to play roles in fine-tuning G protein activation. In plants, non-canonical signaling elements and mechanisms are even more apparent, from the participation of atypical subunits [8, 19, 20], to the paucity of 7TM GPCR-like proteins in plant genomes, of which only GCR1 [21–28] and 7TM1/7TM5 [29] display clear links to G protein signaling. Most strikingly, the role of GPCRs in plants may largely be filled by receptor-like kinases (RLKs) [30].

RLKs comprise the largest family of transmembrane receptors in Arabidopsis [31, 32], and as such are implicated in perceiving a vast array of stimuli including microbial elicitors [33–35], brassinosteroid plant hormones [36], products of cell wall damage [37], phytosulfokines and endogenous danger-associated peptides [38–41], peptides that direct development [42, 43], and peptide attractants that are crucial for fertilization [44]. Subfamilies of RLKs are defined by their extracellular N-terminal domain structure, e.g. leucine-rich repeats (LRR), cysteine-rich, proline-rich, lectin, LysM, wall-associated, malectin-like and S-domain containing subfamilies [45]. Across the broad RLK superfamily, RLKs characteristically contain a single transmembrane spanning domain and a C-terminal kinase domain; however, guanylate cyclase and CaM-binding domains have also been identified within RLK kinase domains, for example, of PSKR1 [46–48] and BRI1 [49, 50].

In Arabidopsis, the first *agb1* mutant was identified in a screen for mutants displaying similarity to mutants of the developmental programming RLK, *ERECTA* [51]. Intriguingly, both *ERECTA*, and *GPA1* are negative regulators of transpiration efficiency, however, epistatic analyses have yet to be reported; in large part due to a strong genetic linkage between the *ERECTA* and *GPA1* loci [52, 53], although this is solvable with newer technologies such as CRISPR. G protein subunits also have been associated with RLKs that perceive the developmentally regulatory class of CLAVATA peptides, such as RPK2 in Arabidopsis [54] and FEA2 in Maize [55]. The functions of the cell death-associated RLKs, BIR1 and SOBIR1, as well as the pathogen-associated RLKs, FLS2, EFR and CERK1, were found to require G protein participation [56] and BIR1, CERK1 and BAK1 were subsequently demonstrated to interact with GPA1, AGG1 and AGG2 [57]. Similarly, Zygote Arrest 1 (ZAR1), an RLK crucial to embryonic development, was found to interact with AGB1 [58]. The role of heterotrimeric G proteins in Arabidopsis pathogen defense responses is well-established [20, 59, 60]; however, mutants of Gβ in Arabidopsis are viable [51, 61], rendering the link to cell-death associated RLKs less clear. Nevertheless, a potential physiological linkage between G proteins and cell death during early development is apparent in other plant species, including rice where Gβ knockout is seedling lethal [62], and maize where Gβ and triple *xlg* knockout mutants are similarly inviable [63, 64]. In both the rice and maize Gβ mutants it was concluded that a runaway autoimmune response was present, based on constitutive expression of defense response genes [62, 64]. Finally, in a previous study we demonstrated that the RLK FERONIA associates with Gβγ, and that a FERONIA ligand, RALF1, stimulates stomatal closure and inhibits stomatal opening in a G protein-dependent manner [65].

The most plausible mechanism by which RLKs signal through heterotrimeric G proteins is by phosphorylation of G protein subunits [66] and their regulatory elements, such as the Regulator of G protein Signaling 1 (RGS1) [67, 68]. In this report we utilized a combination of biochemical, protein-protein interaction and developmental measurements of phosphomimetic mutants to study the effects of phosphorylation at various sites of the canonical Gα protein, GPA1. We found that different phosphosites control distinct aspects of GPA1 function, indicating a specificity of function that allows for a myriad of different phosphorylation-dependent activation and inactivation states beyond the traditional nucleotide-mediated on-off switch of Gα subunits.

## Results

### Identification of potentially impactful GPA1 phosphosites

Multiple sources were queried to identify GPA1 phosphosites with the highest likelihood of resulting in biochemical effects. As we summarized previously [30], the PhosPhat 4.0 database of Arabidopsis phosphosites [69] indicated phosphorylation of a cluster of three N-terminal threonine residues, T12/T15/T19 in published high-throughput datasets [70–75], as well as a G1 GTP-binding site, S49, in an internal dataset that no longer appears in the database, but can still be retrieved from the iPTMnet [76] report page for GPA1. Phosphat 4.0 also annotates a phosphosite on the interdomain cleft between the GPA1 Ras and helical domains, Y166, that has previously been investigated by the Jones group [77, 78]. Subsequently, the Jones group also cataloged numerous *in vitro* phosphorylation events by RLKs on GPA1 that had been preloaded with either GTPγS or GDP [66]. They confirmed that the three N-terminal threonine residues, T12/T15/T19, are phosphorylated by multiple RLKs independently of GPA1 nucleotide status. They also identified additional G1 motif phosphosites, S52 and T53, and a hot spot at an interdomain cleft site, T164, that could be phosphorylated by nine different RLKs [66]. We therefore selected the following subset of phosphosites to study: T12/T15/T19, S49, S52, T53 and T164. We mapped these phosphosites onto high confidence AlphaFold 3 [79] models of GPA1-GDP bound to AGB1/AGG3 (Gγ domain only) and GPA1-GTP bound to RGS1 (Figure 1). The promiscuity of RLKs across the multiple GPA1 phosphorylation sites, as exemplified in the Jia et al. dataset [66], and the poor stability of GPA1 *in vitro* [80] which complicates valid analysis of GPA1 phosphovariants produced by lengthy *in vitro* phosphorylation protocols, dictate that the optimal method for studying the effect of phosphorylation of individual sites both *in vivo* and *in vitro* is by phosphomimetic mutation [81], even if these mutations are not perfectly analogous to native phosphorylation [82].

**Figure 1.**
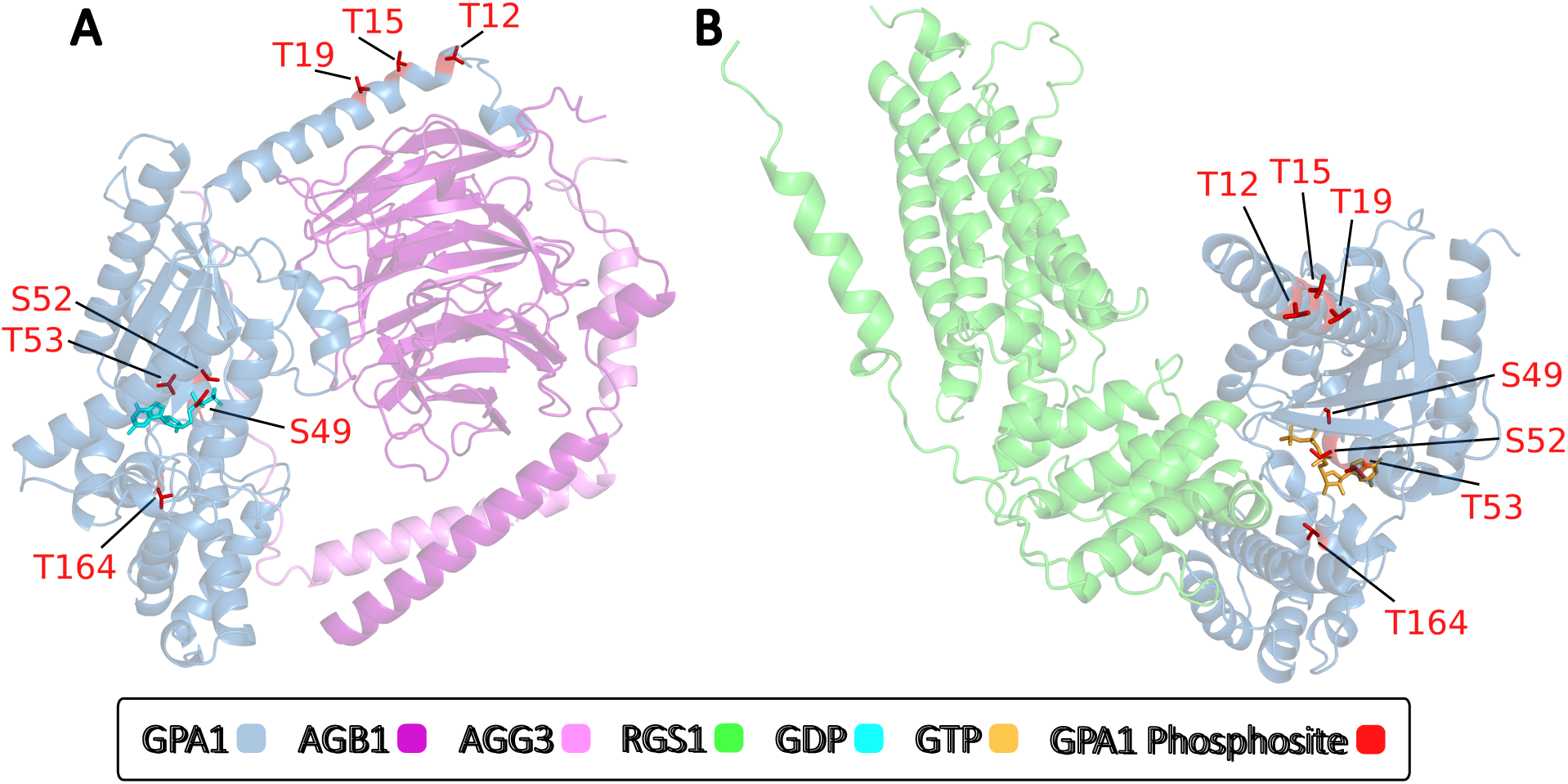
The GPA1 phosphosites assessed are located at the N-terminus or proximal to/within the GPA1 nucleotide binding pocket. T12, T15, T19, S49, S52, T53 and T164 phosphosites of interest were mapped onto high confidence AlphaFold3-generated structural complexes of: **A.** GPA1-GDP-Gβγ3 (piTM confidence score = 0.82) and **B.** GPA1-GTP-RGS1 (piTM = 0.81). Proteins and nucleotides are color coded according to the key, only side chains of the indicated phosphosites are shown, and transparency of the protein backbone was set to 60% for clarity of the positions of the GPA1 phosphosites relative to the nucleotides, Gβγ3 and RGS1. As AGG3 (Gγ3) contains N- and C-terminal extensions that are poorly modelled and not involved in heterotrimer binding, the AGG3 sequence shown corresponds to residues 22-99. For reference, residues 1-112 of AGG3 were used in yeast 3-hybrid assays shown in subsequent figures. All other proteins depicted in Figure 1 are full length.

We utilized three distinct categories of experimental approach for each of the phosphosites: 1) *in vitro* biochemical assays, 2) *in vivo* protein-protein interaction studies, and 3) developmental characterization of lines comprising native promoter-driven complementation of the *gpa1-3* knockout T-DNA mutant line.

### T12/T15/T19 phosphosites

As the T12, T15 and T19 phosphosites are in close proximity and have been identified in the same datasets, we chose to assess phosphonull (T12/15/19A) and phosphomimetic (T12/15/19E) mutation of all three sites simultaneously. These sites on the N-terminal α helix are distal from the nucleotide-binding pocket, so were not expected to substantially alter GTP binding activities, which indeed was the case as assayed by standard G protein activity assays: BODIPY-GTP binding (Figure 2A), the increase in intrinsic tryptophan fluorescence upon the addition of GTPγS (corresponding to the conformational shift of Gα associated with activation (Figure 2B)) and BODIPY-GTPγS and BODIPY-GDP binding (Figure S1). In each of the above assays an increase in fluorescence corresponds to nucleotide binding. Another assay we previously developed [80] assesses the structural stability of GPA1 by SYPRO dye fluorescence in the presence and absence of nucleotide co-factors, in this case GTPγS (time course example shown in Figure S2A). As SYPRO Orange fluorescence increases when bound to hydrophobic regions of proteins, which would normally be buried in the core of the protein, the SYPRO dye acts as a reporter of protein unfolding. Given that GPA1 displays poor stability *in vitro* when not supplemented with additional nucleotides [80], the addition of excess GTPγS strongly stabilizes wild-type GPA1, decreasing SYPRO Orange fluorescence. The SYPRO assay demonstrated similar levels of basal unfolding and GTPγS-stabilization after 15 minutes of incubation at 25°C for wild-type GPA1 and the T12/15/19A and T12/15/19E mutants (Figures S2B and 2C). Together these data indicate that phosphorylation of T12/15/19 sites do not perturb the GDP-GTP cycle of GPA1, a conclusion that is consistent with a recent study by Jia et al. who assayed an S8E/T12E/T15E/T19E GPA1 phosphomimic [78].

**Figure 2.**
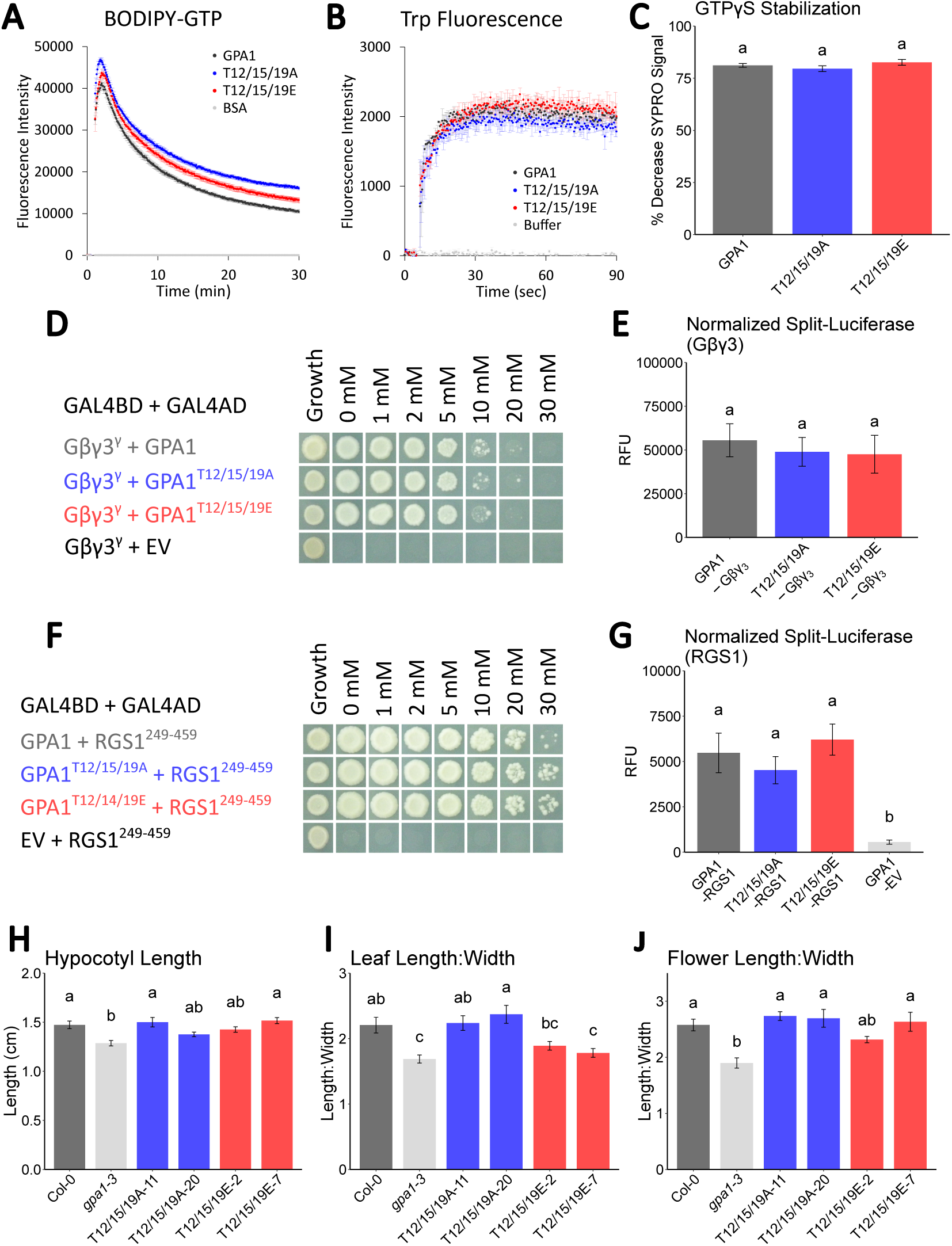
Characterization of GPA1 phosphomutants carrying simultaneous mutation of three N-terminal positions: T12, T15 and T19. **A.** BODIPY-GTP assay of nucleotide binding (increase in signal) and hydrolysis (decrease in signal) activities of GPA1 and T12/15/19A phosphonull and T12/15/19E phosphomimic mutants after injection of the fluorescently labeled BODIPY-GTP nucleotide at T=30 seconds. **B.** Intrinsic tryptophan fluorescence assay reflecting the conformational shift to the active state (increased signal from the Trp residue in switch II) of wild-type GPA1 and GPA1 phosphomutants upon GTPγS injection at T=5 seconds. **C.** The percentage decrease in SYPRO Orange dye fluorescence of 600 nM GPA1 after 15 minutes of incubation with 10 µM GTPγS at 25°C, compared to the minus nucleotide control, indicates the structurally stabilizing effect of GTPγS binding on the GPA1 protein and phosphomutants. **D.** Yeast 3-hybrid protein-protein interaction assay between GPA1 (wild-type and T12/15/19 phosphomutants) and AGG3^1-112^ in the presence of AGB1. **E.** *In planta* split-luciferase assay of wild-type and phosphomutant GPA1L-nFLuc (αB-αC loop tagged GPA1) interactions with cFLuc-AGB1 in the presence of full length AGG3 coexpression. **F.** Yeast 2-hybrid protein-protein interaction assay between GPA1 (wild-type and T12/15/19 phosphomutants) and the cytosolic domain of RGS1, RGS1^249-459^. **G.** *In planta* split-luciferase assay of wild-type and phosphomutant GPA1-nFLuc interactions with RGS1-cFLuc, encoding full length RGS1. **H-J.** Phenotypic characterization of **H.** etiolated hypocotyl length, **I.** leaf roundness (ratio of length:width), and **J.** flower roundness (ratio of length:width) for the *gpa1-3* mutant complemented with native promoter-driven T12/15/19A and T12/15/19E mutants of *GPA1*. See Figure S4 for equivalent assays of wild-type GPA1 and empty vector (EV) controls. In panels **D** and **F**, the “Growth” column corresponds to growth on -Leu-Trp media to confirm yeast viability. The 0-30 mM columns correspond to increasing concentrations of 3-amino-1,2,4-triazole (3-AT), the competitive inhibitor of the HIS3 reporter enzyme, in -Leu-Trp-Met-His interaction selective media. Growth on higher concentrations of 3-AT is interpreted as a stronger, more stable interaction. In panels **C**, **E**, **G**, **H**, **I** and **J,** letters above bars correspond to statistical grouping by one-way ANOVA with the Tukey’s multiple comparison test.

Next we examined the interactions between GPA1 and the AGB1/AGG3 Gβγ dimer that we have shown has the clearest physiological links to GPA1 signaling [19], and between GPA1 and its primary GTPase activating protein, RGS1 [83]. We utilized yeast 3-hybrid and yeast 2-hybrid, as well as *in planta* split-luciferase systems to assay relative interaction strength of wild-type GPA1 vs. phosphomutants of GPA1. As yeast 2-/3-hybrid techniques require the test proteins to be present in yeast cell nuclei, we removed the transmembrane domains, as we previously had, of AGG3 [19] and RGS1 [84]. As no such constraints exist for split-luciferase, we used full length proteins for the *in planta* system. We first established that GPA1 and AGB1 did not autoactivate in the split-luciferase system, by assaying these tagged components in the presence and absence of co-expressed Arabidopsis Gγ subunits. Minimal split-luciferase signal was observed in the absence of Arabidopsis Gγ subunits, while strong positive luminescence signals were observed when each of AGG1, AGG2 and AGG3 were co-expressed (Figure S3A), consistent with our previous BiFC results, which demonstrated Gγ subunits facilitate the interaction between GPA1 and AGB1 *in vivo* [85]. Notably, the GPA1-AGB1 split-luciferase signal was higher in the presence of AGG3 compared to AGG1 or AGG2 (Figure S3A), validating the choice of AGB1/AGG3 as the Gβγ dimer in our protein-protein interaction assays. In the split-luciferase experiments, we utilized a split-firefly luciferase as a readout for interaction and normalized the firefly luciferase data to NanoLuc activity: the GPA1 proteins were tagged with HiBit which could be used to reconstitute the NanoLuc protein in combination with the LgBit protein. As such, NanoLuc activity can also be considered a proxy for GPA1 abundance, which was not statistically different between wild-type GPA1 and the T12/15/19A or T12/15/19E mutants (Figures S3B-C). Our protein-protein interaction assays revealed that no substantial differences from wild-type GPA1 were observed for the interactions with Gβγ3 or RGS1 for the T12/15/19A or T12/15/19E GPA1 mutant proteins in either the yeast or *in planta* systems (Figure 2, D-G).

Finally, we assessed the ability of the GPA1 phosphomutants to complement well-described developmental phenotypes of the *gpa1-3* mutant, specifically: etiolated hypocotyl length, leaf roundness [86] and shorter/wider flowers [87, 88]. The latter two were measured as a ratio of leaf or flower length:width. For all phenotypes, the wild-type GPA1 protein was able to complement the mutant phenotypes and, as expected, the empty vector control construct did not (Figure S4, A-C). While there was some inter-line phenotypic variability observed within the T12/15/19A phosphonull and T12/15/19E phosphomimic lines, all phenotypes were capable of being complemented by both proteins (Figure 2, I-K), except for leaf shape, which was not fully complemented in either T12/15/19E line (Figure 2J).

### S49 phosphosite

The S49 phosphosite resides within the highly conserved G1 nucleotide binding motif, proximal to the β phosphate (Figure 1). The equivalent position has been implicated as a phosphosite in Gα_i2_ after µ opioid receptor stimulation [89] and high-throughput studies have detected phosphorylation at this site of the mammalian Gα_12/13_ family, and the Gα_i_ and/or Gα_s_ families, though phosphorylation events could not unequivocally be mapped to specific Gα proteins due to sequence conservation across Gα subunits of the region around the phosphosite [30]. As expected, the addition of a negative charge in the S49D phosphomimetic mutant all but abolished nucleotide binding (Figures 3A, S5A and B) and as a result nucleotide-mediated conformational change (Figure 3B) and stabilization (Figure 3C). Notably, the level of SYRO Orange fluorescence in the basal state, i.e. in the absence of GTPγS supplementation, was over 10-fold higher than for the wild-type GPA1 protein, indicating substantial accessibility of hydrophobic residues to the dye, consistent with protein unfolding *in vitro*. This raises two possible scenarios that our data cannot unequivocally differentiate between: 1) the S49D mutant is unstable *in vitro* due to a lack of nucleotide binding, or 2) the inherent instability of the S49D mutant *in vitro* results in an unfolded mutant protein. In either scenario the function of the protein is impaired. However, *in vivo*, the S49D mutant protein does appear to fold, as it is capable of interacting with Gβγ3 in yeast (Figure 3D) and *in planta* (Figure 3E), despite notably lower protein accumulation as assessed by NanoLuc signal in the GPA1-Gβγ3 assay (Figure S3B). Strikingly, the S49D mutant is unable to interact with RGS1 (Figures 3F-G), which is consistent with scenario 1 above, in which the S49D mutant fails to bind GTP and therefore does not achieve a transition to the active state, which RGS1 would negatively regulate. Functionally, the S49D phosphomimic mutant is also impaired as it failed to fully complement the *gpa1-3* mutant phenotypes assayed, though it did partially complement the hypocotyl length phenotype. In comparison, the S49A phosphonull mutant displayed wild-type-like structural stability *in vitro* (Figure S2B) and modest reductions in nucleotide binding rates compared to wild-type GPA1 that were much less severe than the loss of nucleotide binding observed for the S49D mutant (Figures 3A-C and S5A-B). S49A did not display any impairment in heterotrimer formation (Figures 3D-E) or RGS1 binding (Figure 3F-G), and was able to fully complement the *gpa1-3* mutant phenotypes (Figure 3H-J). The functional impairment of the phosphomimic compared to the phosphonull suggests a role for phosphorylation of the S49D residue in nucleotide binding/protein stability, RGS1 binding and downstream signaling, with physiological relevance revealed by our developmental measurements.

**Figure 3.**
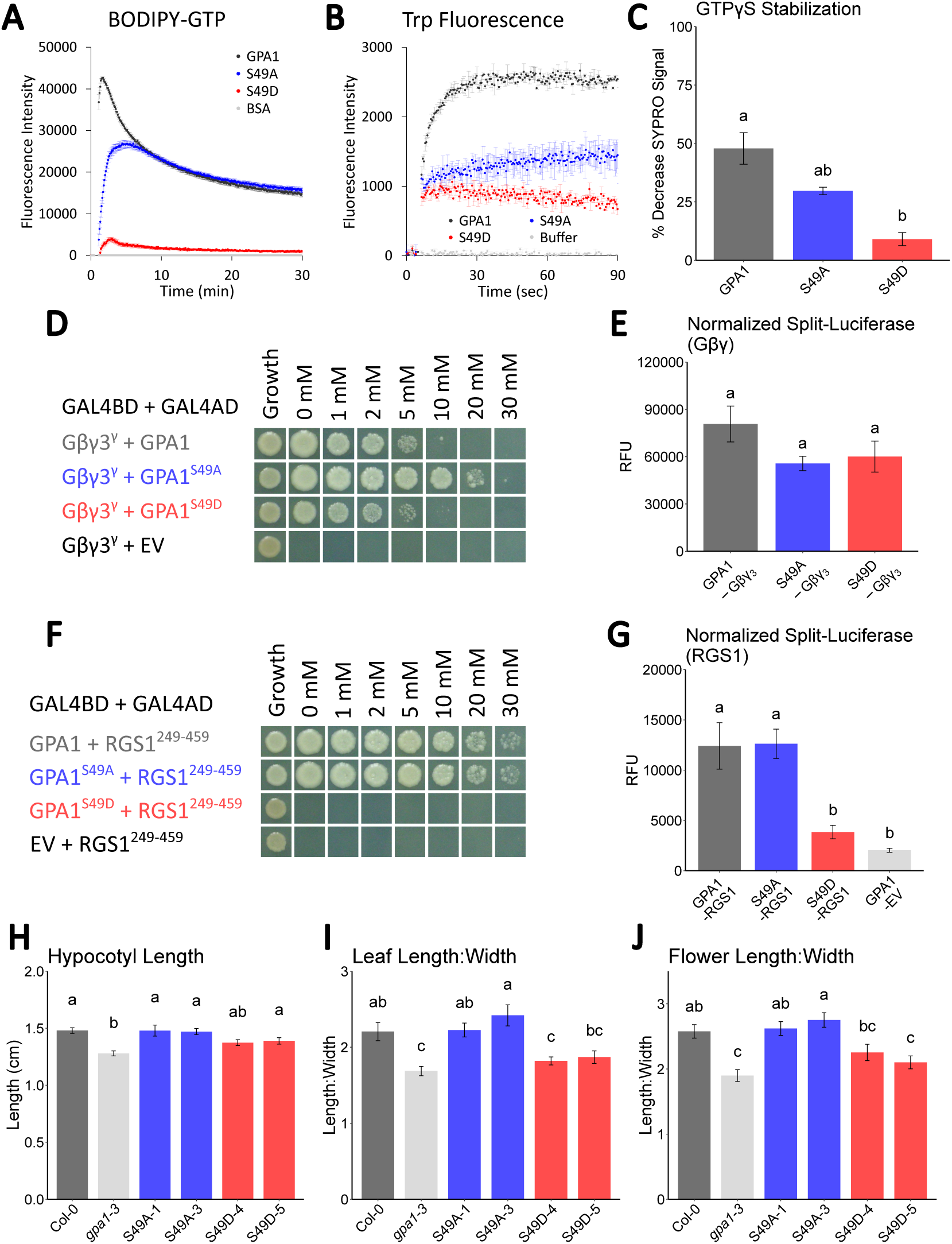
Characterization of GPA1 phosphomutants of the S49 residue within the G1 nucleotide binding motif. **A.** BODIPY-GTP assay of GTP binding and hydrolysis. **B.** Intrinsic tryptophan fluorescence assay with GTPγS added at T=5 seconds. **C.** SYPRO Orange assay of GPA1 stability ±10 µM GTPγS at T=15 minutes. **D** and **E.** Yeast 3-hybrid (**D**) and split-luciferase (**E**) assays of the interactions of wild-type GPA1 and GPA1 phosphomutants with Gβγ3. **F** and **G.** Yeast 2-hybrid (**F**) and split-luciferase (**G**) assays of the interactions of GPA1 and phosphomutants with RGS1. **H-J.** Phenotypic characterization of **H.** etiolated hypocotyl length, **I.** leaf roundness (ratio of length:width), and **J.** flower roundness (ratio of length:width) for the *gpa1-3* mutant complemented with native promoter-driven S49A and S49D mutants of GPA1. Aside from the identity of the phosphomutants of GPA1, all assay details are as described in the Figure 2 legend.

### S52 phosphosite

The first indication that nucleotide binding was not crucial to the function of GPA1 was that an S52C mutant protein was able to complement some *gpa1* mutant phenotypes despite lacking GTP and GDP binding activities [84]. We further showed that an S52N mutant lost GTP binding activity [80], which was expected as this highly conserved position within the G1 is important for coordination of the Mg^2+^ cofactor ion that in turn coordinates GTP binding in both heterotrimeric and small monomeric G proteins [90–92]. As there is evidence that the S52 residue is phosphorylated [66, 93], we investigated the effects of phosphomimic (S52D) and phosphonull (S52A) mutations on GPA1 function. As opposed to previous use of S52C and S52N mutations, we chose to use the S52A mutation as a phophonull for three reasons: 1) the small nature of the alanine side-chain, 2) the hydrophobic alanine side chain (-CH_3_) does not introduce charge or reactivity as S52C or S52N mutations do, and 3) for consistency with the precedent of alanine scanning mutagenesis commonly being used as phosphonull mutants [94]. Both S52A and S52D mutations abolished nucleotide binding (Figures 4A-B and S6A-B) and nucleotide-mediated structural stabilization (Figure 4C) though, unlike for S49D, basal stability of folding *in vitro* was not significantly affected, as shown by basal SYPRO Orange fluorescence (Figure S2B). The S52D mutation strongly impaired the binding of GPA1 to Gβγ3 (Figures 4D-E), while both S52A and S52D mutations impaired binding to RGS1 (Figures 4F-G), consistent with their impaired GTP binding and inability to transition to the active state (Figures 4A-C). The S52C mutant of GPA1 complements *gpa1* mutant hypocotyl, leaf and flower phenotypes [84], which we observed was also the case for S52A (Figures 4H-J). In comparison, the S52D mutant only partially complemented the hypocotyl length phenotype and failed to complement the leaf shape phenotype, although it did complement the flower shape phenotype almost as well as the wild-type protein and the S52A mutant. These results suggest that specifically the phosphomimetic mutation partially impairs cellular function of GPA1, and underscores that the contribution of GPA1 to these three developmental phenotypes is independent of nucleotide binding.

**Figure 4.**
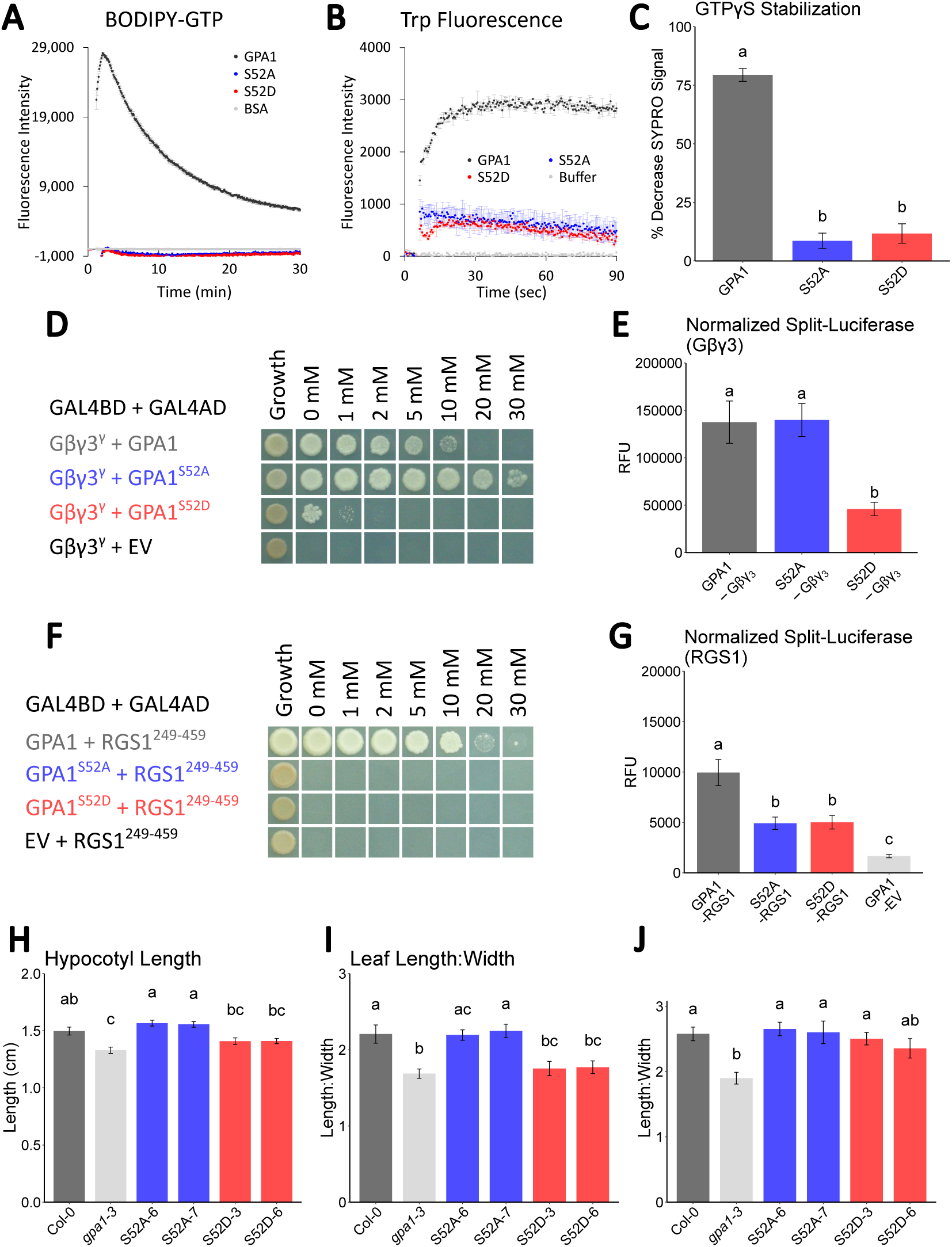
Characterization of GPA1 phosphomutants of the S52 residue within the G1 nucleotide binding motif. **A.** BODIPY-GTP assay of GTP binding and hydrolysis. **B.** Intrinsic tryptophan fluorescence assay with GTPγS added at T=5 seconds. **C.** SYPRO Orange assay of GPA1 stability ±10 µM GTPγS at T=15 minutes. **D** and **E.** Yeast 3-hybrid (**D**) and split-luciferase (**E**) assays of the interactions of wild-type GPA1 and GPA1 phosphomutants with Gβγ3. **F** and **G.** Yeast 2-hybrid (**F**) and split-luciferase (**G**) assays of the interactions of GPA1 and phosphomutants with RGS1. **H-J.** Phenotypic characterization of **H.** etiolated hypocotyl length, **I.** leaf roundness (ratio of length:width), and **J.** flower roundness (ratio of length:width) for the *gpa1-3* mutant complemented with native promoter-driven S52A and S52D mutants of GPA1. Aside from the identity of the phosphomutants of GPA1, all assay details are as described in the Figure 2 legend.

### T53 phosphosite

The final G1 nucleotide binding motif site we investigated was T53, for which evidence of phosphorylation was described in both the Jia et al. [66] and Xue et al. [93] datasets. The T53 site proved to be an interesting case study of potential artifacts introduced by the conjugation of the BODIPY fluorophore to nucleotides when assessing binding pocket mutants. The T53E phosphomimic mutant displayed low or no binding of BODIPY-GTP (Figure 5A), BODIPY-GTPγS (Figure S7A) or BODIPY-GDP (Figure S7B). However, the increase in tryptophan fluorescence upon the addition of GTPγS, indicative of GTPγS binding-induced conformational activation, was not impaired in the T53E mutant (Figure 5B). Similarly, the stabilization of the T53E protein by GTPγS was only slightly, and not statistically significantly, less than that of the wild-type protein (Figure 5C). Furthermore, the T53A mutant displayed slightly higher BODIPY-GTP binding, tryptophan fluorescence in response to GTPγS, and stabilization by GTPγS (Figure 5A-C), but paradoxically, almost no BODIPY-GTPγS binding activity (Figure S7A). The most likely explanation for these discrepancies is that the conjugated BODIPY fluorophore sterically inhibits nucleotide binding in the T53 mutants (Figures 5A and S7), while binding can be observed in assays utilizing unlabelled nucleotides (Figures 5B-C). Interestingly, both T53A and T53E mutants displayed reduced affinity for Gβγ3 (Figures 5D-E) and RGS1 (Figures 5F-G) without a reduction in protein accumulation *in planta* (Figure S3B-C), or loss of stability *in vitro* (Figure S2B). Nevertheless, phenotypic complementation of the *gpa1-3* null mutant was also differential between the mutants, with the T53A mutant complementing all phenotypes, while the T53E mutant almost fully complemented the hypocotyl phenotype (Figure 5H), failed to complement the leaf phenotype (Figure 5I), and partially complemented the flower phenotype (Figure 5J).

**Figure 5.**
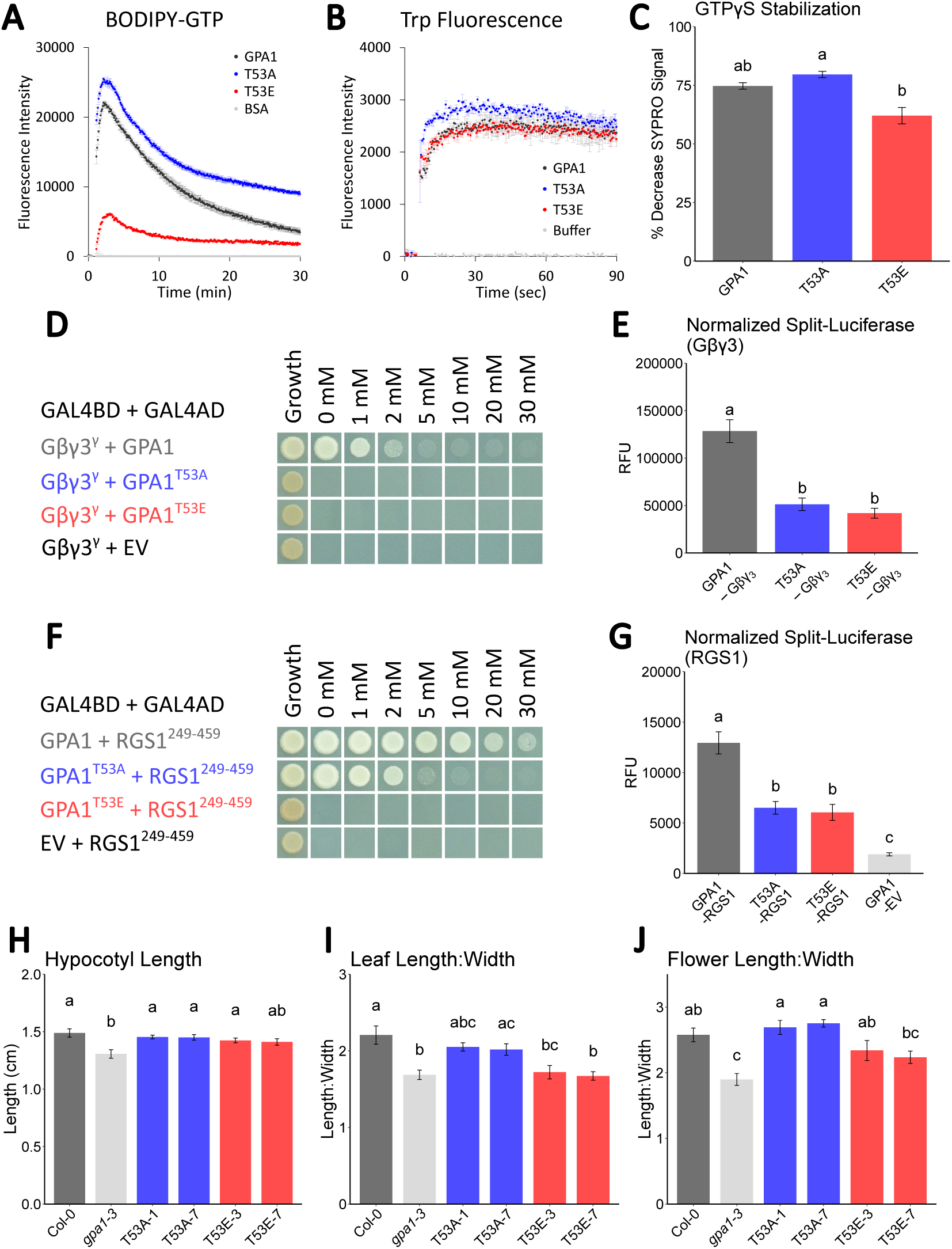
Characterization of GPA1 phosphomutants of the T53 residue within the G1 nucleotide binding motif. **A.** BODIPY-GTP assay of GTP binding and hydrolysis. **B.** Intrinsic tryptophan fluorescence assay with GTPγS added at T=5 seconds. **C.** SYPRO Orange assay of GPA1 stability ±10 µM GTPγS at T=15 minutes. **D** and **E.** Yeast 3-hybrid (**D**) and split-luciferase (**E**) assays of the interactions of wild-type GPA1 and GPA1 phosphomutants with Gβγ3. **F** and **G.** Yeast 2-hybrid (**F**) and split-luciferase (**G**) assays of the interactions of GPA1 and phosphomutants with RGS1. **H-J.** Phenotypic characterization of **H.** etiolated hypocotyl length, **I.** leaf roundness (ratio of length:width), and **J.** flower roundness (ratio of length:width) for the *gpa1-3* mutant complemented with native promoter-driven T53A and T53E mutants of *GPA1.* Aside from the identity of the phosphomutants of GPA1, all assay details are as described in the Figure 2 legend.

### T164 phosphosite

Given the effect of interdomain motion on nucleotide exchange by Gα subunits [14], and that phosphorylation of the interdomain cleft has been shown to enhance the activation of the human Gα subunit, Gα_i3_ [95], we sought to investigate phosphorylation at the T164 position in the interdomain cleft, which was identified to be phosphorylated by nine different RLKs *in vitro* [66]. BODIPY-GTP binding was slightly impaired in the T164E phosphomimic mutant (Figure 6A), while tryptophan fluorescence upon GTPγS binding (Figure 6B) and GTPγS-mediated structural stabilization (Figure 6C) were wild-type-like for both T164A and T164E mutants. Contrastingly, BODIPY-GTPγS binding was abolished in the T164E mutant (Figure S8A) and BODIPY-GDP binding was reduced (Figure S8B). The discrepancy between BODIPY-GTP, tryptophan fluorescence and SYPRO Orange assays (Figures 6A-C) with the BODIPY-GTPγS assay (Figure S8A) most likely again reflects an artifact of BODIPY conjugation, as described above for the T53 mutants. The lack of agreement between BODIPY-GTP and BODIPY-GTPγS assays is a phenomenon we and others have previously observed and attributed to the different positions of conjugation for the BODIPY fluorophore, to the ribose ring in BODIPY-GTP vs. to the γ phosphate in BODIPY-GTPγS [80, 96]. The placement of BODIPY on different ends of GTP may result in steric inhibition of one form, in this case BODIPY-GTPγS, but not always the other. Interestingly, specifically the T164E phosphomimic mutant displayed a reduced affinity for both Gβγ3 and RGS1, while the phosphonull T164A mutant displayed an increased affinity to both in the split-luciferase assay, albeit not statistically significantly so (Figures 6D-G). This opposite effect of the phosphonull and phosphomimic mutations in the split-luciferase system may indicate a portion of the wild-type GPA1 protein is phosphorylated at the T164 position *in planta*. The T164A mutant was also capable of complementing all three phenotypes measured, while the T164E mutant partially complemented the hypocotyl and leaf phenotypes, but fully complemented the flower phenotype (Figure 6H-J).

**Figure 6.**
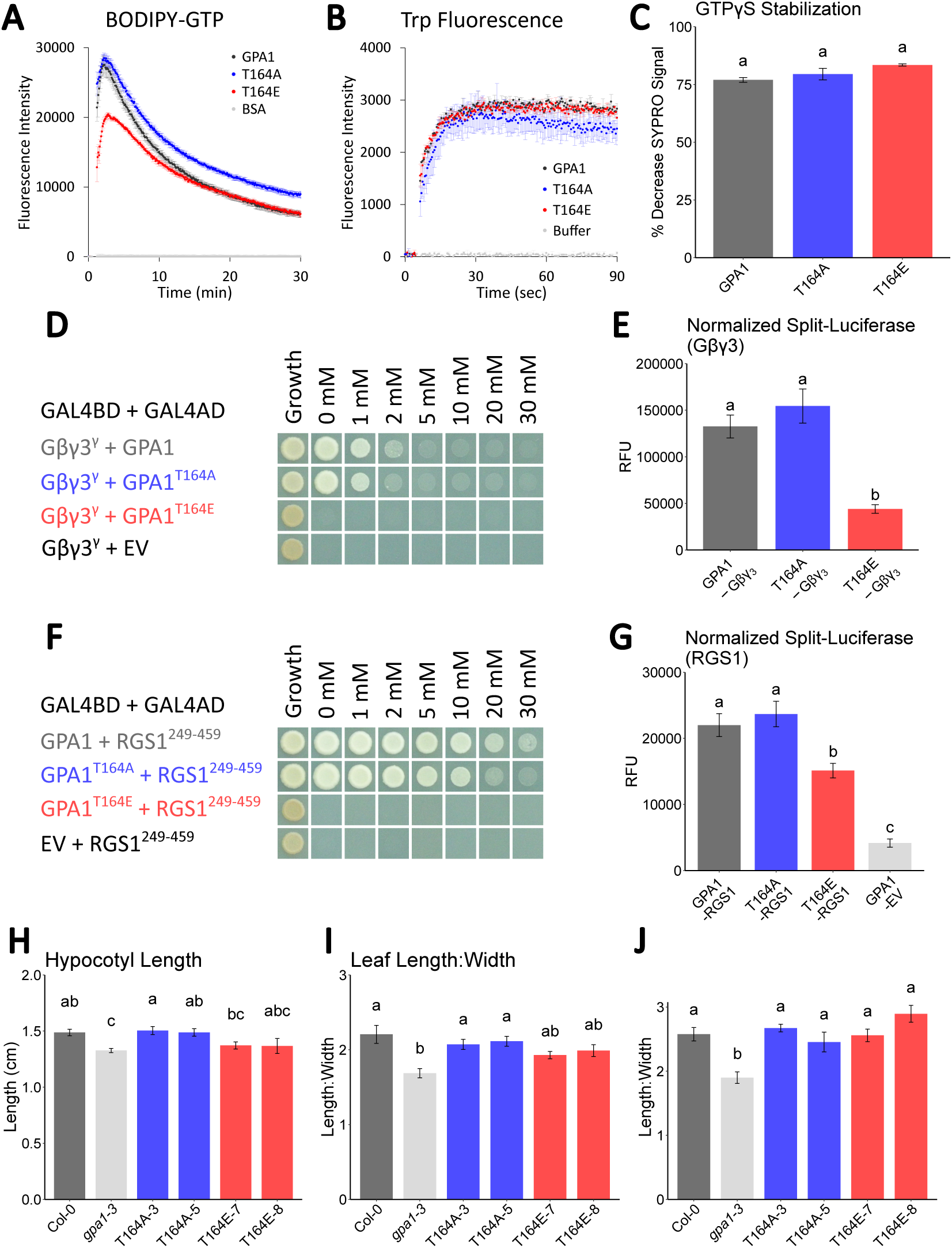
Characterization of GPA1 phosphomutants of the T164 residue within the interdomain cleft. **A.** BODIPY-GTP assay of GTP binding and hydrolysis. **B.** Intrinsic tryptophan fluorescence assay with GTPγS added at T=5 seconds. **C.** SYPRO Orange assay of GPA1 stability ±10 µM GTPγS at T=15 minutes. **D** and **E.** Yeast 3-hybrid (**D**) and split-luciferase (**E**) assays of the interactions of wild-type GPA1 and GPA1 phosphomutants with Gβγ3. **F** and **G.** Yeast 2-hybrid (**F**) and split-luciferase (**G**) assays of the interactions of GPA1 and phosphomutants with RGS1. **H-J.** Phenotypic characterization of **H.** etiolated hypocotyl length, **I.** leaf roundness (ratio of length:width), and **J.** flower roundness (ratio of length:width) for the *gpa1-3* mutant complemented with native promoter-driven T164A and T164E mutants of GPA1. Aside from the identity of the phosphomutants of GPA1, all assay details are as described in the Figure 2 legend.

## Discussion

Our study reveals that not only do plants utilize atypical heterotrimer G protein subunits such as XLGs [4] and type C Gγ subunits [8, 97], but even GPA1, which initially appeared to be a typical Gα subunit, participates in underappreciated signaling mechanisms, namely those involving phosphorylation. Our analysis of GPA1 phosphosites combines assays addressing three different aspects of GPA1 function: nucleotide binding, physical interactions with its cognate Gβγ3 dimer and the GAP protein RGS1, and morphological development. Table 1 compares the effects of each phosphomutant across all of our biochemical, split-luciferase protein-protein interaction, and developmental measurements. The set of GPA1 phosphorylation sites that we studied demonstrates that phosphomimicry alters a diversity of processes differentially, whether it be abolishing GTP binding (S49D and S52D in Figures 3 and 4, respectively), suppressing interactions with Gβγ3 (S52D, T53E and T164E in Figures 4, 5 and 6, respectively) or RGS1 (S49D, S52D, T53E and T164E in Figures 3, 4, 5 and 6, respectively), or impairing GPA1-mediated development (Figures 2-6 and summarized in Table 1). Table 1 reveals that GTP binding activity does not correlate with complementation of hypocotyl, leaf or flower *gpa1* null mutant phenotypes, in agreement with our previous work [84].

**Table 1.** Summary of results for biochemical, protein-protein interaction and developmental measurements for each GPA1 phosphomutant protein.

| Phosphomutant | GTP binding (consensus)* | Interactions** |  | Phenotypic complementation of *gpa1* null mutant |  |  |
| --- | --- | --- | --- | --- | --- | --- |
|  |  | +Gβγ3 | +RGS1 | Hypocotyls | Leaves | Flowers |
| T12/15/19A | WT-like | WT-like | WT-like | Complemented | Complemented | Complemented |
| T12/15/19E | WT-like | WT-like | WT-like | Complemented | *gpa1*-like | Complemented |
| S49A | Reduced | WT-like | WT-like | Complemented | Complemented | Complemented |
| S49D | Abolished | WT-like | Abolished | Partially complemented | *gpa1*-like | *gpa1*-like |
| S52A | Abolished | WT-like | Reduced | Complemented | Complemented | Complemented |
| S52D | Abolished | Reduced | Reduced | Partially complemented | *gpa1*-like | Complemented |
| T53A | WT-like | Reduced | Reduced | Complemented | Complemented | Complemented |
| T53E | WT-like | Reduced | Reduced | Complemented | *gpa1*-like | Partially complemented |
| T164A | WT-like | WT-like | WT-like | Complemented | Complemented | Complemented |
| T164E | WT-like | Reduced | Reduced | Partially complemented | Partially complemented | Complemented |

\* The GTP binding column represents the consensus of the BODIPY-GTP, BODIPY-GTPγS, tryptophan fluorescence and SYPRO Orange stabilization assays, except for T53A, T53E and T164E, where the BODIPY-GTP/-GTPγS results were excluded as we propose that the BODIPY fluorophore introduced artifacts by steric inhibition in these assays.
\*\* The yeast 2-/3-hybrid results largely agree with our split-luciferase data, but where minor discrepancies existed we characterized the interactions based on the split-luciferase assays as, in contrast to the yeast assays, the split-luciferase assays were conducted *in planta* with all full length proteins, including full length AGG3 and RGS1 proteins containing their transmembrane domains.

We propose that phosphorylation at the sites we studied generally plays an antagonistic role in GPA1-mediated development. Mimicking constitutive phosphorylation by expression of T12/15/19E (Figures 2H-I and Table 1), S49D (Figures 3H-I and Table 1), S52D (Figures 4H-I and Table 1), T53E (Figures 5H-I and Table 1) or T164E (Figures 6H-I and Table 1) mutant proteins in the *gpa1-3* background in each case failed to complement at least one of the three phenotypes we examined, indicating a negative regulatory role of each event. However, the mechanism was not universal across tissues for individual phosphosites as specificity of complementation, or lack thereof, was observed between the mutants (Table 1). These distinct effects of different GPA1 phosphosites begin to answer one of the more puzzling questions in plant G protein studies, namely, how so many different signals can be integrated from diverse RLKs through a common heterotrimeric G protein system with a limited repertoire.

In mammalian GPCR signaling, the concept of the phosphorylation barcode in which GPCRs are differentially phosphorylated in order to bias their signaling activity and direct subcellular localization, is well known [98]. We propose that different RLKs each imprint a specific phosphorylation barcode on GPA1, allowing bespoke regulation of the multiple facets of GPA1 signaling in a manner that stimulates the appropriate downstream responses. For example, much as many of the exogenous signals in stomatal development [99] are distinct from those for hypocotyl elongation [100], flower development [101] and pathogen responses [102], the elements downstream of GPA1 are also likely different, and fine tuning of GPA1 signaling to effectors under specific environmental or endogenous conditions could be achieved by differential phosphorylation, allowing for numerous G protein activation/inactivation states.

One mechanistic aspect that could be common to the S49 and S52 phosphosites is the ability to stimulate nucleotide dissociation, specifically receptor-mediated GTP release from GPA1. Receptor-mediated GDP dissociation underpins the canonical activation paradigm of mammalian G proteins. However, it is well-established that GPA1 is a self-activating Gα subunit [80, 103], which potentially minimizes the role of receptor-stimulated GDP release. On the other hand, the recently elucidated phenomenon of GPCR-stimulated GTP release as a mechanism to attenuate active signaling in mammalian cells may be more relevant to plant Gα subunits. In mammals, different ligands of the µ opioid receptor skew the activity of the receptor from promoting GTP loading to favoring GTP release [17, 18]. It could be envisaged that in plants receptor-mediated phosphorylation of GPA1 at sites such as S49 and S52 stimulates GTP unloading to abrogate the signaling of GPA1 in GTP-dependent phenotypes, such as promotion of stomatal development [84]. Though the ligands and classes of receptors have diverged, it is possible that phosphorylation of serines in the G1 motif is an evolutionarily conserved mechanism that attenuates GTP-dependent Gα signaling in mammals and plants.

Comparative analyses of different mutations further underscores the importance of phosphorylation. Though not all of our phosphonull mutants displayed wild-type nucleotide binding activities, they all fully complemented the *gpa1* mutant phenotypes (Table 1). Contrastingly, each of the phosphomimic proteins failed to restore at least phenotype (Table 1), indicating it is not simply non-specific perturbation of conserved residues that impairs GPA1 physiological function, but rather position-specific addition of a negative charge, mimicking phosphorylation, that impairs function. The hypothesized importance of phosphorylation is additionally supported by the ability of the S52C mutant of GPA1 to complement *gpa1* mutant phenotypes [84], much like the S52A mutant (Figures 4H-J), while the S52D mutant was not able to complement hypocotyl or leaf *gpa1* mutant phenotypes (Figures 4H-I). Phosphorylation thus provides a key source of regulation outside of the canonical G cycle.

We selected phosphosites to study based on structural predictions that they were likely to have functional consequences on GPA1 signaling. Consequently, we mostly selected residues within the core of the protein, near or within the nucleotide binding pocket (Figure 1); based on the data summarized in Table 1, the resultant subset of GPA1 phosphomimics were indeed mostly functionally deleterious. Prior publications by Jia et al. [66] and Xue et al. [93] have described additional phosphosites on the surface of GPA1, mostly on the helical domain. The surface phosphorylation events may be involved in direct coupling to receptors or effectors, thereby either promoting or suppressing nucleotide-independent direct signaling activity of GPA1, and are thus worthy of additional study. Indeed, it could be proposed that surface sites are more interesting as they could be more accessible to RLKs and effectors. Nevertheless, access to the binding pocket by kinases is partially explained by the interdomain motion revealed by electron microscopy studies [14]. Furthermore, Xue et al. demonstrated that phosphorylation of the T19 position was necessary for the RLK BAK1 to phosphorylate GPA1, and for flg22-stimulated dissociation of GPA1 from RGS1 [93]. It is also possible that phosphorylation at the N-terminus of GPA1 is a required initial step to open up the protein for phosphorylation in the core. The charge status of the N-terminal α-helix (αN) of Gα subunits also has been described to play a role in GPCR coupling [104, 105], and though 7TM GPCR-like proteins such as GCR1 are not known to play a major role in plant G protein function, we recently confirmed that GPA1 does bind to GCR1 *in vivo* [24], and it is plausible that N-terminal phosphorylation of GPA1 is a mechanism to stimulate 7TM GPCR-Gα complex reconfiguration or dissociation for increased accessibility of GPA1 by RLKs. This potentially explains two phenomena: 1) that the T12/15/19E phosphomimetic mutations had few effects on GPA1 signaling (Figure 2) despite being commonly identified in plant proteomic datasets, and 2) that all 11 of the RLKs that phosphorylated GPA1 in Jia et al.’s phosphoproteomic dataset phosphorylated the T19 position [66], perhaps because this is a necessary step prior to specific phosphorylation at additional sites.

It is plausible that there are additional functionally important phosphosites, including sites that positively regulate GPA1 signaling activity. It is also likely that multiple GPA1 residues are phosphorylated concurrently, and combinatorial complexity is another largely unexplored layer to phosphoregulation of GPA1 as well as mammalian Gα subunits. Indeed these aspects and the critical importance of GPA1 phosphorylation are suggested by a recent report from Jia et al. which demonstrated that an interplay of phosphorylation of the αN and the Y166 interdomain cleft sites promotes the interaction between GPA1 and TCP14/JAZ3 proteins, thus preventing their proteasomal degradation and thereby enhancing immunity against the bacterial pathogen *Pseudomonas syringae* [78]. In our study it is noteworthy that S49D, S52D and T164E phosphomimic proteins individually each partially complement the *gpa1* null mutant hypocotyl phenotype (Table 1), and it is possible that simultaneous phosphomimetic mutation of these sites would more strongly impair GPA1 function in developing hypocotyls.

Finally, the specificity of different phosphomutants in different tissues is perhaps surprising, as despite manifesting at different developmental stages, there is a commonality to the three phenotypes with *gpa1* null mutant hypocotyls, leaves and flowers all sharing shorter and wider phenotypes [86–88]. Despite this, the S49D phosphomimic protein, for example, partially complemented the hypocotyl phenotype, but failed to complement the leaf and flower phenotypes (Figures 3H-I and Table 1). Meanwhile, the S52D phosphomimic protein fully complemented the flower phenotype while showing no or partial ability to complement the other phenotypes (Figures 4H-I and Table 1), and the T53E mutant was not impaired in restoring the hypocotyl phenotype, but was unable to fully complement the leaf and flower phenotypes (Figures 5H-I and Table 1). These results indicate context-specific signaling roles for each phosphorylation event that await additional elucidation.

## Methods

### Plant growth

All transgenic Arabidopsis complementation lines were generated in the *gpa1-3* mutant (ABRC stock CS6533) background, which is a SALK T-DNA knockout in the Col-0 background [106]. Seeds were surface sterilized, spread on plates containing 0.5X Murashige and Skoog (MS) medium, 1% sucrose and 1% agar (Sigma, St. Louis, MO, USA), and stratified at 4°C in the dark for 2 days. After 10–12 days’ growth on vertical plates in a growth chamber with an 8 hour light/16 hour dark cycle, a light intensity of 220 µmol m^-2^ s^-1^ and a temperature of 21±0.5°C, uniform seedlings were transferred from agar plates to individual 8 x 8 x 9 cm pots containing Pro-Mix BX Mycorrhizae soil (Premier Tech Growers and Consumers Inc.) and returned to the same growth conditions for 6 weeks before leaf blade dimensions and shape were scored. Plants were then shifted to a 16 hour light/8 hour dark cycle to promote flowering for measurement of flower dimensions as previously described [8]. Note: all control and test lines were grown together in a randomized configuration, so the Col-0 and *gpa1-3* values are shared between all figures for leaf and flower shape measurements, i.e. panels I and J in each of Figures 2-6. Hypocotyl assays were conducted on etiolated seedlings growing on 0.5X Murashige and Skoog (MS) medium, 1% sucrose and 1% agar (Sigma, St. Louis, MO, USA) plates, as previously described [87].

### Cloning

Primer sequences used in the cloning steps described in this section are provided in Table S1. A pCR8/GW/TOPO clone containing a complementary DNA (cDNA) copy of the *GPA1* open reading frame (pCR8-GPA1) was used as a template for downstream tagging and mutagenesis. A hemagglutinin (HA) epitope tag sequence was added to the αB-αC loop of GPA1 (encoded amino acids 131 and 132) by overlap extension PCR, and the tagged *GPA1L-HA* sequence was TA cloned into pCR8/GW/TOPO (pCR8-GPA1L-HA). Phosphomimic (aspartic acid [D] or glutamic acid [E]) and phosphonull (alanine [A]) mutations were introduced into pCR8-GPA1 or pCR8-GPA1L-HA sequences by overlap extension PCR (T12/15/19A, T12/15/19E, S49A, S49D, S52A and S52D) or REPLACR mutagenesis (T53A, T53E, T164A and T164E) [107].

Wild-type and mutant sequences were amplified from pCR8-GPA1 clones with 5’ NcoI and 3’ Kpn2I sites adapted for cloning into the protein expression vector, pSTTa that encodes N-terminal dual StrepII-tags for purification, as we previously described [80]. *GPA1* sequences were also mobilized from pCR8-GPA1 clones into pDEST-GADT7 and pDEST-GBKT7 yeast 2-hybrid vectors by Gateway LR reaction. pBridge constructs for expression of Arabidopsis G□γ for yeast 3-hybrid were previously described [19]. The fragment of *RGS1* encoding the cytosolic RGS-containing domain (residues 249-459) was amplified from Arabidopsis seedling cDNA and TA cloned into pCR8/GW/TOPO. The sequence encoding the RGS1 cytosolic fragment was then mobilized into pDEST-GADT7. We adapted pDOE vectors [85] for four different tag configurations (N- vs. C-terminal tagging of each protein: pSLU-01→04) with split-firefly luciferase (FLuc) components and a C-terminal HiBit tag on the nFLuc fusion for normalization. For split-luciferase experiments, KflI and AatII sites were adapted by PCR to the *AGB1* coding region for cloning into MCS3 of pSLU-02. The coding region of *GPA1* was first cloned into the NcoI-SpeI sites of MCS1 in pSLU-04 to generate an nFLuc-GPA1 sequence with a C-terminal HiBit tag. Overlap extension PCR was used to introduce the N-terminus of firefly luciferase into the αB-αC loop of GPA1 (same position as the HA tag above), a 5’ NcoI site and the 3’ end of GPA1 was amplified from pSLU-04-GPA1 to include the C-terminal HiBit tag and 3’ XbaI site. This GPA1L-nFLuc-HiBit fragment was cloned into MCS1 of the above pSLU-02 vector containing *AGB1*, between NcoI/XbaI sites, such that it replaced the existing nFLuc and HiBit sequences. pCR8-GPA1 clones carrying mutations were used as templates for overlap extension fragments to generate αB-αC loop-nFLuc fusions, as above, which were cloned into NcoI-XbaI sites for mutations prior to residue 131, or NcoI-SpeI sites for mutations after position 131, of the above pSLU-02-AGB1 vector. These resulted in vectors encoding GPA1L-nFLuc and cFLuc-AGB1 test constructs, with wild-type and mutant GPA1 sequences, recapitulating the tag configuration that was previously validated for GPA1-G□γ interactions assayed by BiFC [85]. For coexpression of Arabidopsis Gγ subunits, coding regions of AGG1, AGG2 or AGG3 were mobilized from pCR8/GW/TOPO into the pEarleyGate201 [108] plant transformation/expression vector by Gateway LR reaction. The generation of a pSLU-01 vector with an *RGS1* ORF in MCS3 that had been mutagenized to remove a native NcoI site was previously described [24]. Sequences encoding mutated *GPA1* ORFs were cloned into the NcoI/SpeI restriction sites of pSLU-01-RGS1 MCS1, as was previously described for the wild-type *GPA1* sequence [24], yielding test constructs encoding C-terminally tagged GPA1-nFLuc-HiBit and RGS1-cFLuc. Native promoter GPA1 complementation constructs were generated by first amplifying a fragment of the *GPA1* locus including a 3827 bp *GPA1* promoter/5′ UTR in addition to 34 bp after the start of translation, from Col-0 genomic DNA. A cDNA copy of the *GPA1* coding region was also amplified and both fragments were simultaneously ligated into the pORE-O3 binary vector using an ApaI restriction site 5′ of the promoter and a NotI restriction site 3′ of the coding region, with a native PstI site within the 5′ 17 bp of the coding region, that is common to both fragments, used to seamlessly join the promoter/5′ UTR fragment to the coding region and trim redundant sequence between the amplified fragments. A 481 bp *GPA1* 3′ UTR/terminator fragment was then adapted to the 3′ end of the GPA1 coding region by overlap extension PCR, which also introduced a Bsu36I site by mutation of the sequence immediately 3′ of the *GPA1* stop codon from CCTTATT to CCTTAGG. This partial coding region with adapted 3′ UTR/terminator fragment was cloned into the vector between the above internal PstI restriction site and a 3′ NcoI site in the vector. Wild-type and mutated *GPA1* fragments from pCR8-GPA1L-HA cDNA clones were amplified and used to replace the equivalent sequence of the GPA1 coding region in the pORE-O3-GPA1 vector between a naturally occurring *GPA1* SalI site located 23-28 bp 3′ of the start of translation, and the introduced Bsu36I site. Constructs were sequence-verified and then used for Agrobacterium-mediated floral dip transformation [109] of the *gpa1-3* mutant. T3 or T4 homozygous lines were used for developmental measurements.

### Biochemical assays

Wild-type and mutant variants of GPA1 were expressed in high salt LB medium and purified using tandem StrepII-tags as previously described [80]. BODIPY assays of nucleotide binding and SYPRO Orange assays of protein stability were performed as previously described [80]. BODIPY-GDP assays (BODIPY FL GDP, ThermoFisher #G22360) were conducted utilizing the same protocol as the BODIPY-GTP assays. In this report, the percentage decrease of SYPRO Orange signal ±10 µM GTPγS after 15 minutes of incubation at 25°C was plotted as a readout of the structurally stabilizing effect imparted on GPA1 by GTPγS binding. Intrinsic tryptophan fluorescence [110, 111] was assayed in a plate reader format. Proteins were diluted with EB base (25 mM Tris-HCl, 50 mM NaCl and 5% glycerol pH 7.4) to 420 nM supplemented with 5.25 mM MgCl_2_. Ninety-five µl of each sample in a FLUOTRAC 200 96 well half area plate (Greiner Bio-One, 675076) was loaded into a Synergy Neo2 plate reader (Biotek) and baseline intrinsic tryptophan fluorescence was measured in well mode with a 280/10 nm excitation and 340/10 nm emission for 5 seconds with a 0.5 second kinetic interval. Reactions were initiated by injection of 5 µl of 40 µM GTPγS, yielding a final reaction volume of 100 µl containing 400 nM GPA1, 5 mM MgCl_2_ and 2 µM GTPγS. Intrinsic tryptophan fluorescence was monitored as above for an additional 90 seconds. To mitigate the effects of the previously described temperature-dependent instability of GPA1 [80], reactions were only performed for two technical replicates of an individual sample within a single plate reader run, before the next sample was loaded on the plate and assayed. Note: the wild-type GPA1 and buffer control values are shared between Figures 4B and 6B, as tryptophan fluorescence assays of S52 (Figure 4) and T164 (Figure 6) mutants were conducted at the same time with batches of recombinant protein prepared in parallel.

### Protein-protein interaction assays

Yeast 3-hybrid assays between GPA1 and G□γ subunits and yeast 2-hybrid assays between GPA1 and the cytosolic domain of RGS1 were conducted as previously described [19, 84]. Note: some yeast 2-/3-hybrid interaction tests were performed simultaneously for multiple phosphomutants, and as a result the yeast spots corresponding to wild-type GPA1 + G□γ3 or wild-type GPA1 + RGS1 positive controls, as well as empty vector-containing negative control combinations are shared between Figures 5D and 6D, as well as 5F and 6F, specifically between T53 and T164 assays. Split-luciferase assays were conducted as previously described [24]; in the case of heterotrimer assays pSLU-02-GPA1L-nFLuc–cFLuc-AGB1 constructs (OD600 = 0.2) were co-agroinfiltrated with pEarleyGate201-AGG1, -AGG2 or -AGG3 constructs (OD600 = 0.4). Raw firefly luciferase values were normalized to NanoLuc fluorescence, which estimates GPA1 expression, via the abundance of the HiBit tag, as previously established [84].

## Acknowledgments

We thank David Arginteanu for technical assistance. Research reported in this publication was supported by the National Institute of General Medical Sciences of the National Institutes of Health under award number R35GM153492, with prior support from award number R01GM126079.

