## Supplemental materials for "Phosphovariants of the canonical heterotrimeric Gα protein, GPA1, differentially affect G protein activity and Arabidopsis development"

This file includes:

Figure S1
Figure S2
Figure S3
Figure S4
Figure S5
Figure S6
Figure S7
Figure S8
Table S1

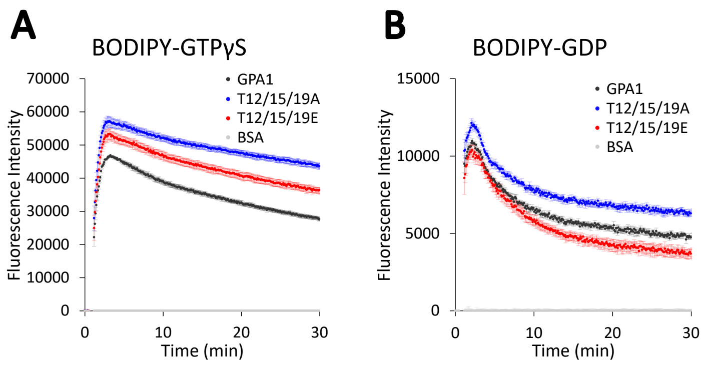


**Figure S1. Additional data for Figure 2. A.** BODIPY-GTPγS and **B.** BODIPY-GDP assays of nucleotide binding for wild-type GPA1 and GPA1 T12/15/19 phosphomutants. Assays were conducted essentially as for BODIPY-GTP in Figure 2, with the fluorescent BODIPY labeled nucleotides injected at T=30 seconds and binding (increased fluorescence) monitored for 30 minutes.

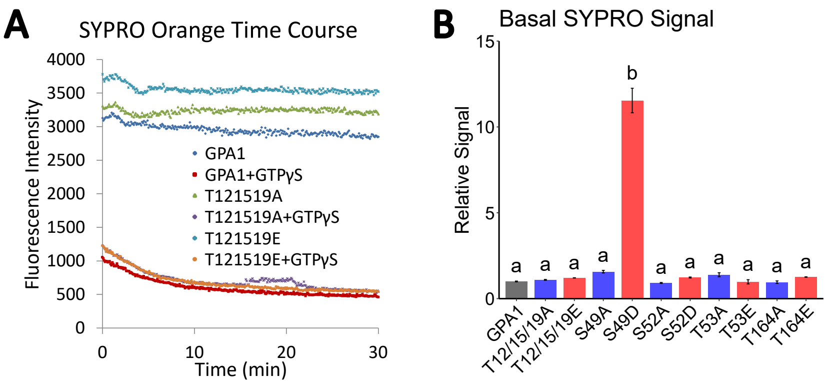


**Figure S2. SYPRO Orange dye fluorescence control data. A.** An example time course assay of 600 nM GPA1 and T12/15/19 triple phosphomutants incubated for 30 minutes at 25 °C ±10 µM GTPγS, during which time fluorescence was monitored. In the main text figures, the reduction in SYPRO fluorescence at T=15 minutes, compared to the respective control lacking nucleotide supplementation, is expressed as a percentage of the control to illustrate the stabilizing effect of nucleotide binding on each mutant. **B.** Basal SYPRO signal corresponds to the average SYPRO Orange signal at T=15 minutes in the non-nucleotide supplemented samples, which was normalized to the wild-type GPA1 control within each assay for comparison of all phosphomutants assayed across multiple days. Aside from the S49D mutant, most GPA1 proteins were similarly folded and stable *in vitro*, regardless of their ability to bind nucleotides.

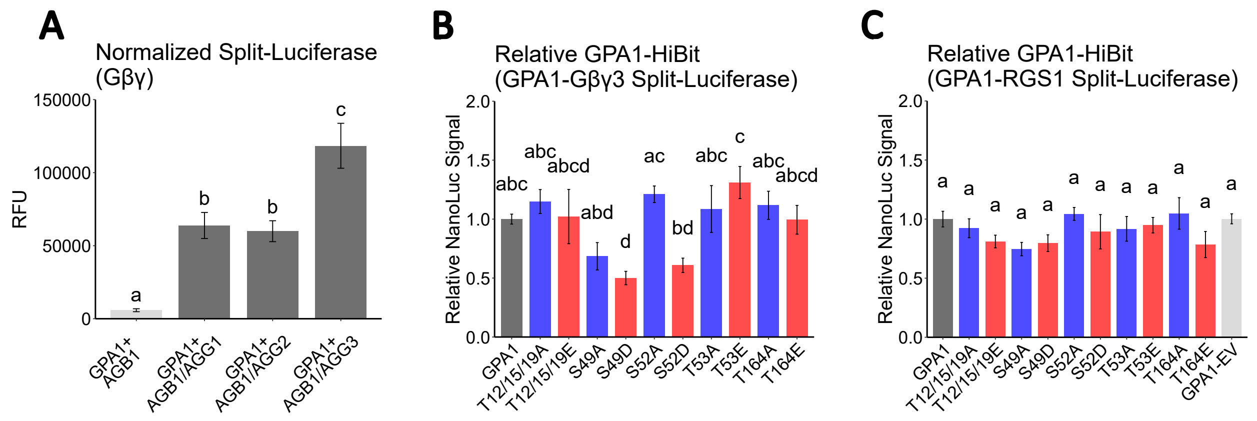


**Figure S3. Split-luciferase control data. A.** Control assay of GPA1L-nFLuc + cFluc-AGB1 demonstrating the specificity of the luminescence signal. Only in the presence of supplemental Gγ expression (AGG1, AGG2 or AGG3), was strong positive signal, indicative of an interaction, observed. **B** and **C.** Relative values (compared to the wild-type GPA1 protein) of control NanoLuc luminescence values against which the split-luciferase (split FLuc) values were normalized. In each assay configuration (GPA1+Gβγ3 in panel **B** and GPA1+RGS1 in panel **C**), the HiBit fragment of NanoLuc was fused to the C-terminus of GPA1, allowing for relative quantification of protein abundance based on NanoLuc signal after reconstitution of the HiBit tag with the exogenously provided LgBit fragment of NanoLuc.

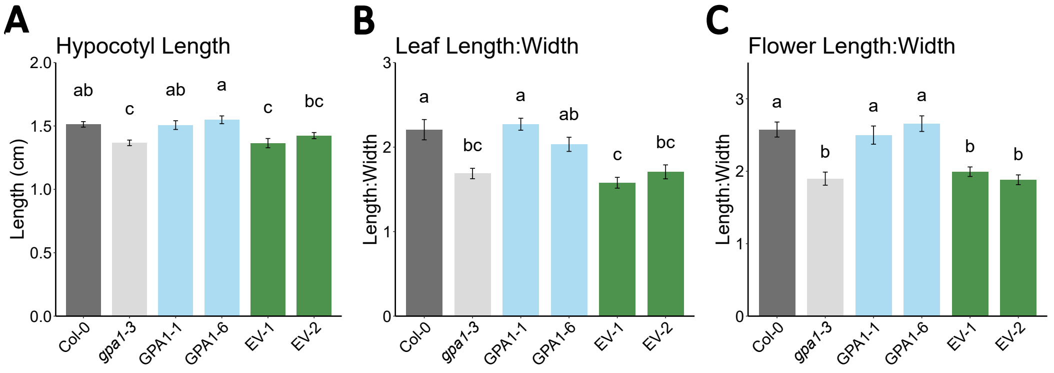


**Figure S4. Control data for complementation of the *gpa1-3* mutant with wild-type GPA1 or empty vector negative controls. A.** Etiolated hypocotyl length, **B.** the ratio of leaf length:width and **C.** the ratio of flower length:width are presented as controls for the complementation of the *gpa1-3* with phosphomutants as displayed in panels H-J of Figures 2-6.

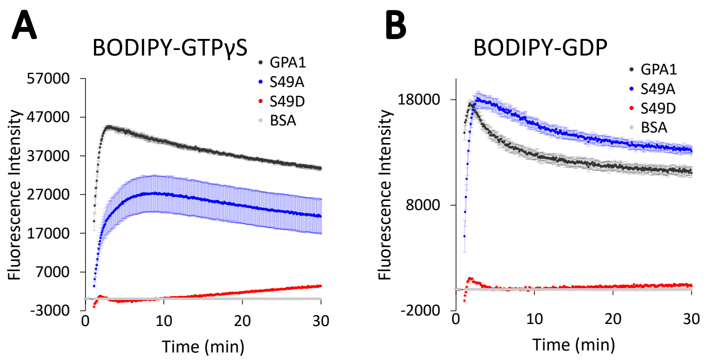


**Figure S5. Additional data for Figure 3. A.** BODIPY-GTPγS and **B.** BODIPY-GDP assays of nucleotide binding for wild-type GPA1 and GPA1 S49 phosphomutants. Assays were conducted essentially as for BODIPY-GTP in Figure 3, with the fluorescent BODIPY labeled nucleotides injected at T=30 seconds and binding (increased fluorescence) monitored for 30 minutes.

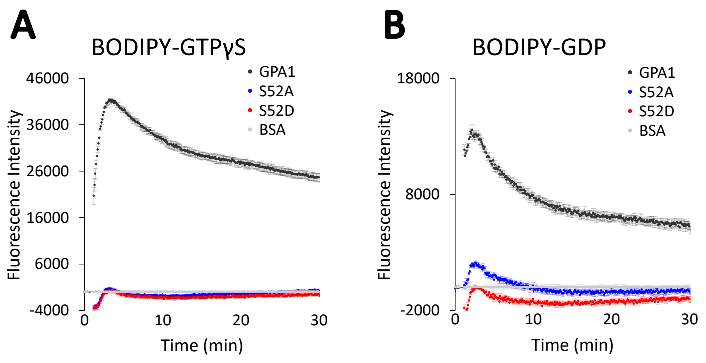


**Figure S6. Additional data for Figure 4. A.** BODIPY-GTPγS and **B.** BODIPY-GDP assays of nucleotide binding for wild-type GPA1 and GPA1 S52 phosphomutants. Assays were conducted essentially as for BODIPY-GTP in Figure 4, with the fluorescent BODIPY labeled nucleotides injected at T=30 seconds and binding (increased fluorescence) monitored for 30 minutes.

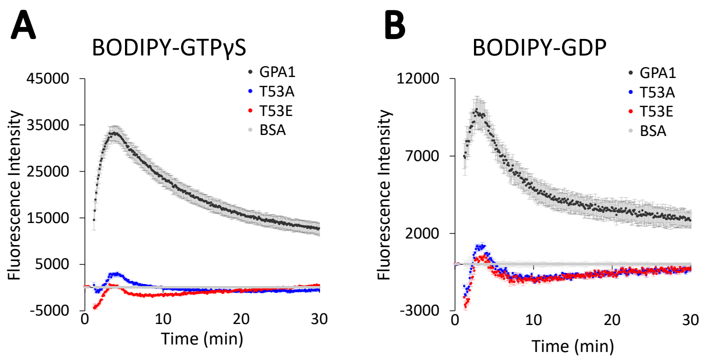


**Figure S7. Additional data for Figure 5. A.** BODIPY-GTPγS and **B.** BODIPY-GDP assays of nucleotide binding for wild-type GPA1 and GPA1 T53 phosphomutants. Assays were conducted essentially as for BODIPY-GTP in Figure 5, with the fluorescent BODIPY labeled nucleotides injected at T=30 seconds and binding (increased fluorescence) monitored for 30 minutes.

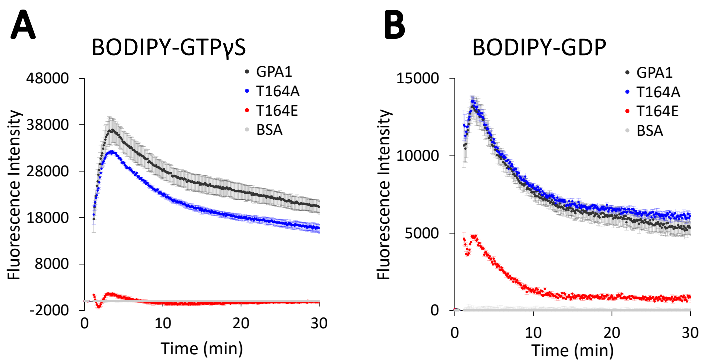


**Figure S8. Additional data for Figure 6. A.** BODIPY-GTPγS and **B.** BODIPY-GDP assays of nucleotide binding for wild-type GPA1 and GPA1 T164 phosphomutants. Assays were conducted essentially as for BODIPY-GTP in Figure 6, with the fluorescent BODIPY labeled nucleotides injected at T=30 seconds and binding (increased fluorescence) monitored for 30 minutes.

**Table S1.** Sequences and descriptions of primers used in this study.

| Primer | Sequence | Use |
|---|---|---|
| GPA1f | ATGGGCTTACTCTGCAGTA | Cloning *GPA1* into pCR8 |
| GPA1rs | TCATAAAAGGCCAGCCTCC | |
| AGG1f | ATGCGAGAGGAAACTGTGG | Cloning *AGG1* into pCR8 |
| AGG1rs | TCAAAGTATTAAGCATCTGCA | |
| AGG2f | ATGGAAGCGGGTAGCTCC | Cloning *AGG2* into pCR8 |
| AGG2rs | TCAAAGAATGGAGCAGCCA | |
| AGG3f | ATGTCTGCTCCTTCTGGC | Cloning *AGG3* into pCR8 |
| AGG3rs | TTAGAAAGCTAAACAACAAGG | |
| RGS1F1 | ATGAACCAACCTCTTCTTTCACA | Cloning the cytosolic domain of *RGS1* into pCR8 |
| RGS1rs | TTAACCGGGACTACTGCATC | |
| GPA1-T12/15/19E-of | GAGGAAGATGAGGATGAGAATGAGCAGGCTGCTGAAATCG | Overlap extension PCR for the T12/15/19E mutant of *GPA1* |
| GPA1-T12/15/19E-or | CTCATTCTCATCCTCATCTTCCTCATGATGTCGACTTCTACTGCAGAG | |
| GPA1-T12/15/19A-of | GCAGAAGATGCAGATGAGAATGCACAGGCTGCTG | Overlap extension PCR for the T12/15/19A mutant of *GPA1* |
| GPA1-T12/15/19A-or | TGCATTCTCATCTGCATCTTCTGCATGATGTCGACTTCTACTGCAGAG | |
| GPA1S49D-of | GGTGCTGGGGAAGACGGAAAATCTAC | Overlap extension PCR for the S49D mutant of *GPA1* |
| GPA1S49D-or | GTAGATTTTCCGTCTTCCCCAGCACC | |
| GPA1S49A-of | GGTGCTGGGGAAGCAGGAAAATCTAC | Overlap extension PCR for the S49A mutant of *GPA1* |
| GPA1S49A-or | GTAGATTTTCCTGCTTCCCCAGCACC | |
| GPA1S52D-of | ATCTGGAAAAGACACAATTTTTAAGCAGA | Overlap extension PCR for the S52D mutant of *GPA1* |
| GPA1S52D-or | GCTTAAAAATTGTGTCTTTTCCAGATTCC | |
| GPA1S52A-of | ATCTGGAAAAGCAACAATTTTTAAGCAGA | Overlap extension PCR for the S52A mutant of *GPA1* |
| GPA1S52A-or | GCTTAAAAATTGTTGCTTTTCCAGATTCC | |
| GPA1-T53E-rf | GAAAATCTGAAATTTTTAAGCAGATAAAACTTCTATTCC | REPLACR mutagenesis for the T53E mutant of *GPA1* |
| GPA1-T53E-rr | TGCTTAAAAATTTCAGATTTTCCAGATTCCCCAGC | |
| GPA1-T53A-rf | AAAATCTGCAATTTTTAAGCAGATAAAACTTC | REPLACR mutagenesis for the T53A mutant of *GPA1* |
| GPA1-T53A-rr | GCTTAAAAATTGCAGATTTTCCAGATTCCCCAGC | |

| Primer | Sequence | Use |
|---|---|---|
| GPA1-T164E-rf | TGATTGTGAGAAATATCTGATGGAGAACTTGAAGAG | REPLACR mutagenesis for the T164E mutant of *GPA1* |
| GPA1-T164E-rr | GATATTTCTCACAATCAGGAACCTGAAGCTCA | |
| GPA1-T164A-rf | TGATTGTGCGAAATATCTGATGGAGAAC | REPLACR mutagenesis for the T164A mutant of *GPA1* |
| GPA1-T164A-rr | GATATTTCGCACAATCAGGAACCTGAAGC | |
| GPA1L-HA-of | TACCCATACGATGTTCCAGATTACGCTACCAAGGACATCGCTGAG | Overlap extension PCR for internal HA tagging of *GPA1* |
| GPA1L-HA-or | AGCGTAATCTGGAACATCGTATGGGTAAAGACGTGGATAGTCTAACC | |
| GPA1f-NcoI | GGGCCATGGGCTTACTCTGCAG | Cloning *GPA1* into pSTTa |
| GPA1r-BspEI | GGGTCCGGATCATAAAAGGCCAGCCTCCAG | |
| AGB1f-KflI | GCCGGGTCCCTAATGTCTGTCTCCGAGCT | Cloning *AGB1* into MCS3 of pSLU-02 to generate a cFLuc-AGB1 fusion |
| AGB1rs-AatII | GGCGACGTCTCAAATCACTCTCCTGTGTC | |
| RGS1-NcoI-rf | TGTTGCCTTGGATCTTCGGAGC | REPLACR mutagenesis to synonymously edit an *RGS1* NcoI site |
| RGS1-NcoI-rr | GAAGATCCAAGGCAACAAAACAAGGG | |
| RGS1 RsrII 34F | ATGCGGTCCGGATGTGCTCTACATGGTGGTTGTC | Cloning *RGS1* into MCS3 of pSLU-01 to generate an RGS1-cFLuc fusion |
| RGS1 AatII 1355R | TCGGACGTCCCGGGACTACTGCATCTGGAAC | |
| GPA1f-NcoI | GGGCCATGGGCTTACTCTGCAG | Cloning *GPA1* into MCS1 of pSLU vectors |
| GPA1rns-SpeI | GGGACTAGTTAAAAGGCCAGCCTCCAG | |
| GPA1-NLuc-or | TCCTGACCCACCTCCGGAGGTAAGACGTGGATAGTCTAACCTACC | Overlap extension PCR for internal nFLuc tagging of *GPA1* |
| Link-NLuc-of | TCCGGAGGTGGGTCAGGAATGGAAGACGCCAAAAACATAAAGAAAGG | |
| NLuc-loop-or | GGATCCTGACCCACCTCCAC | |
| GPA1-NLuc-of | GTGGAGGTGGGTCAGGATCCAAGGACATCGCTGAGGGAATAG | |
| HiBitr-XbaI | GGGTCTAGATTAGCTAATCTTCTTGAACAG | Cloning already HiBit-tagged products into pSLU vectors |
| GPA1 ug-3802 ApaI F | ATCGAGGGCCCTTTCAACTTCTTCTTTTGTGTTATTTGG | Forward primer for *GPA1* promoter (pORE-O3 cloning) |
| GPA 5R | TATGATGTCGACTTCTACTGCAGAGTAAGC | Reverse primer at 5' end of the *GPA1* CDS (pORE-O3 cloning) |
| GPA 29F +ATG | ATGGGCTTACTCTGCAGTAGAAGTCGACA | Forward primer for GPA1 *CDS* (pORE-O3 cloning) |
| GPA1 NotI Rs | CTGGAGGCTGGCCTTTTATGAGCGGCCGCTTT | Reverse GPA1 *CDS* primer with NotI site (pORE-O3 cloning) |

| Primer | Sequence | Use |
|---|---|---|
| GPA1 ~LL 1120Rs | GACACTAAGAAGGAGAAATCTCTTGGAGGCTGGCCTTTTATGA | Reverse *GPA1* CDS primer for overlap extension PCR to join the *GPA1* terminator (pORE-O3 cloning) |
| GPA1 ~LL Bsu36I F | GAAATCTCTTGGAGGCTGGCCTTTTATGACCTTAGGATTACATATCTCTA | Forward *GPA1* terminator primer for overlap extension PCR to join the *GPA1* CDS (pORE-O3 cloning) |
| GPA1 digR NcoI | CAGAACATTCGAAAAAGCGACTTTACCATGGAGAGA | Reverse primer for *GPA1* terminator |
| GPA1r-Bsu36I | CCCCCTAAGGTCATAAAAGGCCAGCCTCCAG | Reverse *GPA1* CDS primer with Bsu36I site (pORE-O3 cloning) |
